# Current-voltage characteristics of K+ channels estimated by MD simulations and Markov State Models

**DOI:** 10.64898/2026.02.04.703779

**Authors:** Simone Furini, Luigi Catacuzzeno

## Abstract

Molecular dynamics (MD) simulations have yielded important insights into ion conduction in potassium channels, but quantitative comparison with electrophysiological experiments remains challenging. Due to their high computational cost, MD simulations are typically performed at membrane potentials well above physiological values, and at only a limited number of voltages. Since current-voltage relationships are not necessarily linear, this limits direct comparison between simulations and experiments. Here, we introduce a method to estimate the current-voltage characteristics of ion channels from Markov state models (MSMs) constructed from MD simulations performed at only a few membrane potentials. Time-discrete MSMs of ion conduction are converted into continuous-time rate matrices, whose voltage dependence is modelled using a rate theory formulation with free energy barriers depending on membrane potential. This approach enables the prediction of channel currents over a wide voltage range without additional simulations. We validated the method using MD simulations of the potassium channels KcsA and MthK. In both cases, the currents predicted at low membrane potentials are in good agreement with those obtained directly from MD simulations, demonstrating the robustness and efficiency of the approach.

## Introduction

Potassium channels are integral membrane proteins that allow the selective transmembrane permeation of potassium ions along an electrochemical gradient. Molecular Dynamics (MD) simulations based on the experimentally determined structures have significantly contributed to the current understanding of conduction mechanisms in these ion channels (Furini & Domene, 2013)(Roux, 2017). The region responsible for the selective conduction of potassium ions, known as the Selectivity Filter (SF), comprises five binding sites for K^+^, designated S0 to S4, starting from the extracellular entrance (Figure 1) (Zhou *et al*., 2001). At these binding sites, a fully or partially dehydrated potassium ion is surrounded by oxygen atoms from the channel protein. Based on atomistic simulations, two mechanisms were proposed for ion conduction through the SF, the soft knock-on and the hard knock-on, which differ in the presence or absence of water in-between the permeating ions. In soft knock-on, single files of potassium ions alternated by water molecules move in concert along the SF (Aqvist & Luzhkov, 2000)(Morais-Cabral *et al*., 2001)(Bernèche & Roux, 2001). In hard knock-on, water molecules do not participate in ion conduction, and potassium ions occupy adjacent binding sites (S2 and S3) at the core of the SF (Furini & Domene, 2009)(Köpfer *et al*., 2014).

**Figure 1.**
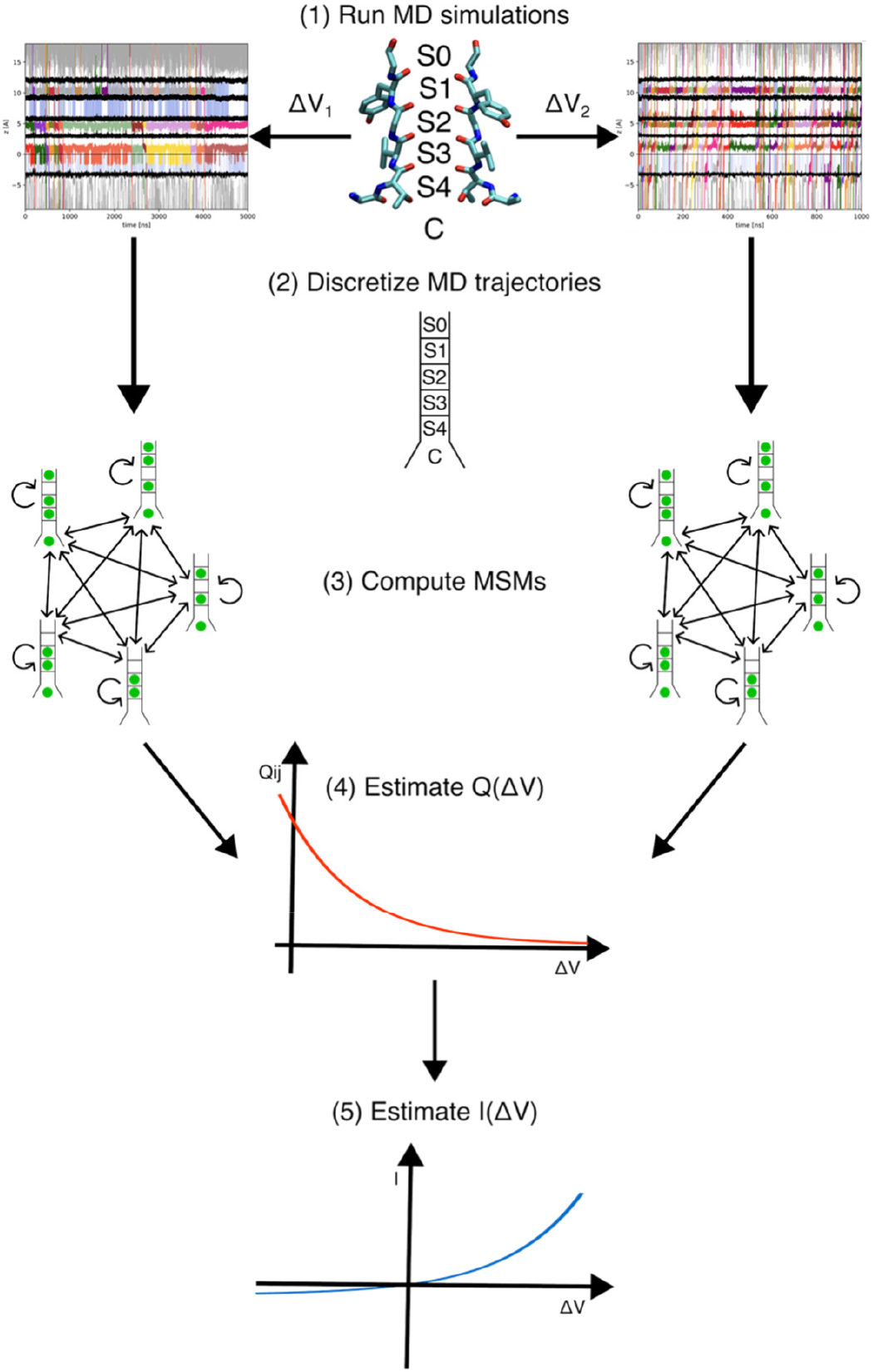
Workflow diagram. The algorithm includes the following steps: (1) Simulate conduction events at different membrane potentials, here indicated as ΔV_1_ and ΔV_1_; (2) Discretize the MD trajectories; (3) Compute MSMs of conduction events; (4) Estimate the voltage-dependence of the rate constants; and (5) Use the voltage-dependent rate matrixes to compute the current-voltage characteristic. In step (1), the atomic structure of the SF is shown for two opposing subunits in licorice representation, together with labels for binding sites S0-S4, and the intracellular cavity, C. The snapshots from the MD trajectories are discretized using the ion occupancy of these binding and cavity, as schematically depicted in step 2. In the schematic representation of the SF green circles are used for binding sites occupied by potassium ions.

An effective strategy for analysing conduction in ion channels at the atomic scale is to directly reproduce the ion movements in MD simulations in the presence of a drifting force acting across the channel, for instance, by applying an electric field perpendicular to the lipid membrane (Gumbart *et al*., 2012), and then to use Markov State Models (MSMs) for extracting pertinent information from these MD trajectories (Harrigan *et al*., 2017)(Furini & Domene, 2018). One advantage of MSMs is their ability to be estimated from independent MD trajectories, and to provide a simplified, easily understandable, description of permeation events. Besides, once an MSM is defined, it provides a theoretically robust framework for analysing conduction and comparing different experimental conditions. MSMs have been employed to analyse how the conduction properties of potassium channels depend on the specific ion channel characteristics, force field, or boundary conditions (Domene *et al*., 2021)(Kei & de Groot, 2023).

An important feature of MSMs estimated from MD simulations is that they can be used, instead of the original MD trajectories, to predict the experimental properties of molecular systems. In the case of ion channels, the most common and meaningful comparison between MSMs predictions and experimental data concerns channel conductance. To facilitate these calculations, we recently developed a method to calculate ion currents directly from MSMs of conduction events (Catacuzzeno *et al*., 2024).

This ability to calculate ion currents directly from MSMs facilitates the comparison with experimental data. However, when considering the comparison between MD simulations and electrophysiological experiments, it is crucial to highlight two important issues. Firstly, in order to accelerate sampling, MD simulations are typically performed at membrane potentials higher than those adopted in experiments. Membrane potentials as high as 500/750 mV are not uncommon in the literature, whereas in experiments they rarely exceed 100-150 mV. Since neither the experimental nor the estimated conductances are guaranteed to exhibit a linear dependence with voltage, comparing conductance values obtained at different membrane potentials could introduce significant inaccuracies and biases. This observation is directly related to the second issue that commonly affects the comparison between simulated currents and experiments. Due to computational constraints, MD simulations are usually limited to a single, rarely a few, membrane potentials. Instead, to assess whether MD simulations can reproduce the functional properties of an ion channel derived experimentally, it would be more informative to compare current-voltage characteristics.

To address these limitations, we developed a method to estimate the current–voltage characteristics of ion channels using MSMs derived from MD simulations performed at only a limited number of membrane potentials. A schematic overview of the workflow is shown in Figure 1. In summary, MD simulations carried out at high membrane potentials are first used to construct time-discrete MSMs of ion conduction, which are subsequently converted into continuous-time rate matrices. The voltage dependence of the transition rates is then modelled using exponential functions based on a rate theory formulation, in which the activation free energies depend on the membrane potential. The resulting voltage-dependent rate matrices are finally used to estimate channel currents, as well as other dynamical properties of ion conduction, over a range of membrane potentials.

The method was tested on two prototypical potassium channels, KcsA and MthK, simulated using different force fields. In both cases, the currents and conduction properties predicted at low membrane potentials from MSMs estimated from MD simulations at high membrane potentials were in good agreement with those computed directly from MD simulations at the same membrane potentials.

## Methods

### Molecular Dynamics (MD) Simulations

The MthK model is based on the Protein Data Bank entry 3LDC, representing the channel in an open/conductive state (Ye *et al*., 2010). The model includes residues from Val18 to Ile99 corresponding to the transmembrane region of the protein. Residues His68 and Cys77 were mutated respectively to Ser and Val, to match the amino acid sequence of the wild-type protein. The channel was inserted in a pure POPC lipid membrane and solvated in an 800 mM KCl solution of TIP3 water molecules (∼15.000 molecules).

The model of the KcsA channel is based on the experimental structure of the non-inactivating E71A mutant in the open/conductive state, Protein Data Bank entry 5VK6 (Cuello *et al*., 2017). The entire transmembrane domain of the channel, from residue Trp26 to residue Gln121, was considered. The lipid membrane is a mixture of 1-palmitoyl-2-oleoyl-glycero-3-phosphocholine (POPC) and 1-palmitoyl-2-oleoyl-sn-glycero-3-phosphate (POPA), in a 3:1 POPC/POPA ratio. The system was solvated with TIP3P water molecules (*∼* 15,000 molecules) and ions, to reach a final concentration of 200 mM KCl.

The initial systems were built with CHARMM-GUI (Wu *et al*., 2014) using the Orientations of Proteins in Membranes (OPM) database to place the channels in the lipid membranes (Lomize *et al*., 2012). Potassium ions were manually placed at binding sites S0, S2, and S4. Atomic models were initially equilibrated by 10,000 steps of energy minimization, followed by 10 ns of dynamics in the NPT ensemble with a 1 fs timestep and 60 ns in the NPT ensemble with a 2 fs timestep. During the equilibration phase, restraints on protein and lipid atoms were gradually reduced to zero. The simulations with the external electric field were performed in the NVT ensemble. Membrane potentials were obtained by applying constant electric fields acting in the direction perpendicular to the lipid membrane (Gumbart *et al*., 2012).

Simulations of the MthK model were executed with GROMACS2023 (Abraham *et al*., 2015). The CHARMM36m (Huang *et al*., 2016) force field was used, with charge scaling factor equal to 0.75, and the corresponding adjustments for the vdW ion parameters (Hui *et al*., 2025). vdW interactions were truncated at 12 Å, and smoothly switched-off in the range 10-12 Å. Long-range electrostatic interactions were calculated with the particle mesh Ewald method using a grid spacing of 1.2 Å and a cutoff of 12 Å for long-range interactions (Essmann *et al*., 1995). The LINCS algorithm was used to restrain bonds with hydrogen atoms (Hess *et al*., 1997). The temperature was set at 310 K and controlled by velocity rescaling with a stochastic term (Bussi *et al*., 2007). In NPT simulations, the pressure was set to 1 atm by semi-isotropic coupling with time constant equal to 5 ps (Bernetti & Bussi, 2020). Trajectories were dumped with a 2 ps time period.

Simulations of the KcsA model were executed using NAMD2 (Phillips *et al*., 2005). The ff14sb version of the AMBER force field was used (Maier *et al*., 2015) in combination with ion parameters by Joung and Cheatham for the TIP3P water model (Joung & Cheatham, 2008) (Jorgensen *et al*., 1983). vdW interactions were truncated at 9 Å. Standard AMBER scaling of 1−4 interactions was applied. Long-range electrostatic interactions were calculated with the particle mesh Ewald method using a grid spacing of 1.0 Å (Essmann *et al*., 1995). The SETTLE algorithm was used to restrain bonds with hydrogen atoms (Tuckerman *et al*., 1992). The temperature was set at 310 K and controlled by coupling the system to a Langevin thermostat with a damping coefficient of 1 ps^−1^. In NPT simulations, a pressure of 1 atm was maintained by coupling the system to a Nose−Hoover Langevin piston, with a damping constant of 25 ps and a period of 50 ps (Feller *et al*., 1995). Trajectories were dumped with a 10 ps time period. Tables S1 and S2 in Supplementary Material show the entire set of simulations with external electric field, respectively for the MthK and KcsA model systems.

### Discretization of the MD trajectories

Trajectories were analysed using the python library MDanalysis and the SciPy ecosystem (Michaud-Agrawal *et al*., 2011)(Virtanen *et al*., 2020). Visual Molecular Dynamics (VMD) was used to inspect trajectories and to generate images of the systems (Humphrey *et al*., 1996). The centres of mass of the following set of atoms were considered as boundaries for the different channel regions, respectively in the KcsA/MthK models: residues Ala107/Glu92 (intracellular boundary of the channel cavity), hydroxyl oxygen atom of residues Thr75/Thr59 (intracellular boundary of S4), backbone oxygen atom of residues Thr75/Thr59 (intracellular boundary of S3), backbone oxygen atoms of residues Val76/Val60 (intracellular boundary of S2), backbone oxygen atom of residues Gly77/Gly61 (intracellular boundary of S1), backbone oxygen atom of residues Tyr78/Tyr62 (intracellular boundary of S0), and backbone oxygen atom of residues Gly79/Gly63 (extracellular boundary of S0). Ions were considered in binding sites S0-S4 or in the cavity when they were located between the boundaries of the corresponding regions (Figure 1) and within 4 Å from the channel axis for binding sites S0 to S4, or 8 Å for ions in the cavity. The occupancy of the 5 binding sites, S0 to S4, and of the channel cavity by potassium ions was used to convert the simulated trajectories into sequences of discrete states, that were then used to estimate the Markov State Models.

### Markov’s State Models (MSMs) of ion conduction

Time discrete MSMs are defined by the following equation:

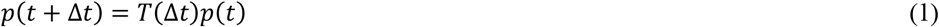

Where *p*(*t*) is a column vector representing the probability of the system being in any state at time *t*, and *T*(Δ*t*) is the transition matrix estimated with sampling period Δ*t*, with element *T*_*ij*_(Δ*t*) representing the probability to move from state *j* to state *i* in Δ*t*. These probabilities were estimated by counting the transitions between states in the discretized MD trajectories:

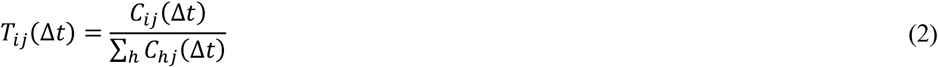

Where *C*_*ij*_(Δ*t*)is the number of transitions from state *j* to state *i* in the MD trajectories sampled with period Δ*t*. The eigenvector of the transition matrix corresponding to the eigenvalue with norm equal to 1 gives the stationary distribution of the system among microstates. This stationary distribution, *π*, was used to identify the set of microstates above a predefined threshold, *π*_*low*_(set to 10^-2^). The procedure was repeated for each membrane potential used for fitting the voltage-dependence of the rate constants, as described in the next sections. Then, the set of microstates with stationary probability above *π*_*low*_ at any fitting membrane potential was identified, and all the MSMs were projected onto this set of states, 𝒮. The projection was performed by merging any microstate that was not part of 𝒮 to the state belonging to𝒮 with maximum transition probability;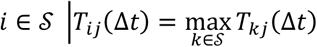. The transition matrix in the new set of states, *T*^𝒮^ (Δ*t*), was computed using the following equations based on conservation laws:

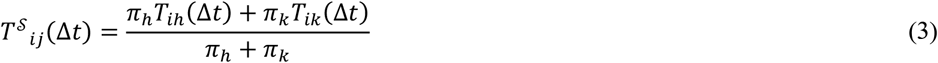

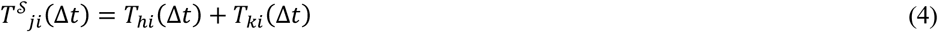

Where *h* and *k* are the microstates to be merged into state *j*.

### Time-continuous MSMs of ion conduction

In the time-continuous formulation of Markov’s model, the analogous of equation (1) is:

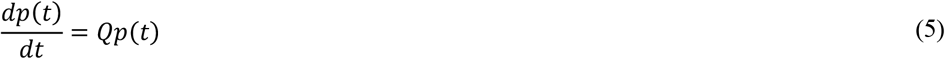

where *Q* is the rate matrix of the system. The transition matrix and the rate matrix are related by the equation:

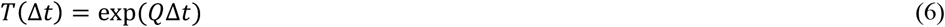

With exp being the matrix exponential function.

In case of an MSM that describes ion conduction across an ion channel, once the rate matrix *Q* is defined, the channel current can be computed as:

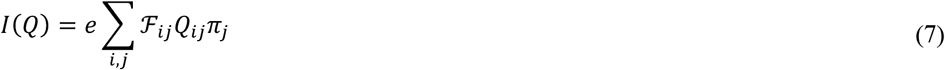

Where ℯis the elementary charge; and ℱ_*ij*_ is the fraction of elementary charge that moves from the intracellular to the extracellular side of the channel for the transition from state *j* to state *i*. The elements of the Flux Matrix, ℱ_*ij*_, were defined considering the number of ions that moves from the intracellular to the extracellular side of the channel along the pathway that requires the minimum number of ion movements (Catacuzzeno *et al*., 2024).

The rate matrix *Q* for the MSM with set of states 𝒮 was computed by maximizing the function:

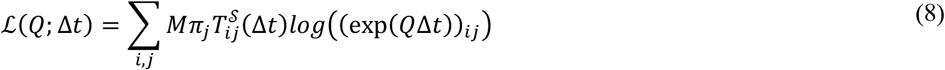

Where *M* is the total number of samples used to estimate the MSM, *log* represents the scalar logarithmic function, and exp is the matrix exponential function, as in equation (6). Intuitively, equation (8) measures how likely is that the states transitions observed in MD trajectories are generated from a time-continuous MSM with rate matrix 𝒬. Only the out-of-diagonal elements of *Q* were estimated numerically by maximizing equation (8). The diagonal elements were computed as *Q*_*ii*_ = − ∑_*h≠i*_ *Q*_*hi*_ to guarantee formal consistency of the rate matrix. The numerical maximization of equation (8) was performed under these constraints: 1) the current estimated by equation (7) deviates less than 5% from the current computed directly by counting the number of conduction events in MD trajectories; 2) the out of diagonal elements of *Q* are positive. The reason for imposing the first constraint is that the inverse of the matrix exponential function, equation (6), is not unique; it is consequently necessary to limit the searching space to solutions that are compatible with the channel current observed in MD trajectories. A step-by-step strategy was used to estimate the rate matrix consistent with the transition matrix at sampling time Δ*t*. Firstly, the rate matrix corresponding to the transition matrix at the shortest sampling time Δ*t*_*O*_ was computed, using as initial conditions for the out of diagonal elements 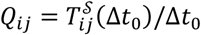. Then, the rate matrixes at increasing sampling times were estimated using as initial conditions at each sampling time the corresponding estimates of the out of diagonal elements of *Q* at the previous sampling time. The SciPy implementation of the algorithm SLSQP (Sequential Least SQuares Programming) was used for the numerical optimizations (Virtanen *et al*., 2020).

### Voltage Dependent MSMs of ion conduction

The effect of the membrane potential, Δ*V*, on the rate constants, *Q*_*ij*_ with *i* ≠ *j*, was modelled with an exponential function (Woodhull, 1973):

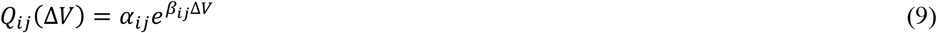

The parameters *α*_*ij*_ and *β*_*ij*_ were estimated by minimizing the mean square distance to the rate constants estimated from MD data. This fitting procedure can lead to numerical instabilities when the rate constants used for fitting are close to zero. This is due to the fact that these low rate constants are affected by large uncertainties, as the corresponding transitions are rarely sampled in MD trajectories, and small changes in *Q*_*ij*_ might cause wide differences in the estimated parameters *α*_*ij*_ and *β*_*ij*_. To correct for these numerical instabilities, the following procedures were applied. Firstly, in case of an *i, j* element that was estimated from rate constants that are below a pre-defined threshold (set to 10^-5^) at all the membrane potentials used for fitting, *α*_*ij*_ was set to zero. This is motivated by the fact that the rate constant for this transition is practically zero at all the simulated membrane potentials, suggesting that it does not contribute significantly to the channel dynamics. A more complicated situation appears when the rate constant for the transition from *j* to *i* is significantly different from zero only at one of the fitting membrane potentials. In this case, it is not possible to exclude a significant contribution of this transition at some membrane potentials, but at the same time, the fitting of the parameters *α*_*ij*_ and *β*_*ij*_ with the procedure described above is clearly affected by large uncertainties. These critical parameters *α*_*ij*_ and *β*_*ij*_ were refined by minimizing the mean square distance to all the rate constants in column *j*, including the diagonal element *Q*_*jj*_. In this way, information from the other rate constants on the same column of the rate matrix are shared, which improves the accuracy in the estimates of these critical parameters. Once all the *α*_*ij*_ and *β*_*ij*_ are defined, the rate matrix can be computed at any membrane voltage, and, in combination with the ℱ matrix it can be used to estimate channel current at membrane potentials not used in the fitting procedure.

### Uncertainty Estimates

To estimate the uncertainty affecting all the parameters, the simulated trajectories were divided in blocks of 50.000 snapshots each, which corresponds to 500 ns and 100 ns of simulated time for the KcsA and MthK channel, respectively. Then, 20 datasets were generated by bootstrapping with resampling among these blocks. In both channels, the length of the blocks is significantly longer than the single conduction events. All values in the Results section are reported as averages and standard deviations computed by repeating the analyses in these 20 datasets generated by bootstrapping.

## Results

### MthK channel

The current voltage characteristic of the MthK model was estimated using MD simulations with membrane potentials equal to -250 mV and +250 mV, respectively. The sampling time for estimating the MSMs was set to 0.1 ns, in accordance with the observations that the timescales of the MSMs are practically independent of the sampling time for sampling time above this value (Figure S1). At steady-state, only 7 states have probabilities above 0.01 at -250 mV or +250 mV (Table S3). Both MSMs, at -250 mV and +250 mV, were projected onto this reduced set of 7 states using conservation laws as defined in equations (3) and (4). These 7 states represent more than 0.98 of the cumulative probability (0.983 at -250 mV, and 0.994 at +250 mV). Therefore, the projection of the MSMs to this reduced set of states is not expected to have a severe impact on the accuracy of the predictions, while it highly facilitates the numerical estimation of the rate matrixes.

The numerical procedure used to estimate the time-continuous MSMs returned rate matrixes that, once converted into transition matrixes by equation (6), are in almost perfect agreement with the transition matrixes estimated by MD trajectories at all the sampling times (Figure S2).

The voltage-dependence of the rate constants was modelled by an exponential function, equation (9), using as input data only the rate constants computed by MD at membrane potential equal to -250 mV and +250 mV, respectively. The entire rate matrix as a function of the membrane potential is reported as supplementary material (Figure S3), while two representative cases are shown in Figure 2. The transition from a state with ions in C, S4, S3, S1 to a state with ions in C, S4, S2, S1 has a strong voltage dependence, which is well-reproduce by an exponential function (Figure 2A). Instead, the voltage-dependence is almost null for the transition from a state with ions in C, S4, S3, S1 to a state with ions in C, S4, S3, S0 (Figure 2B). The state transitions in Figure 2A and 2B corresponds to the movement of one ion from S3 to S2 and from S1 to S0, respectively. Thus, the strong voltage-dependence of the first rate constant, and not of the second one, agrees with the idea that most of the voltage-drop occurs within the core of the SF.

**Figure 2.**
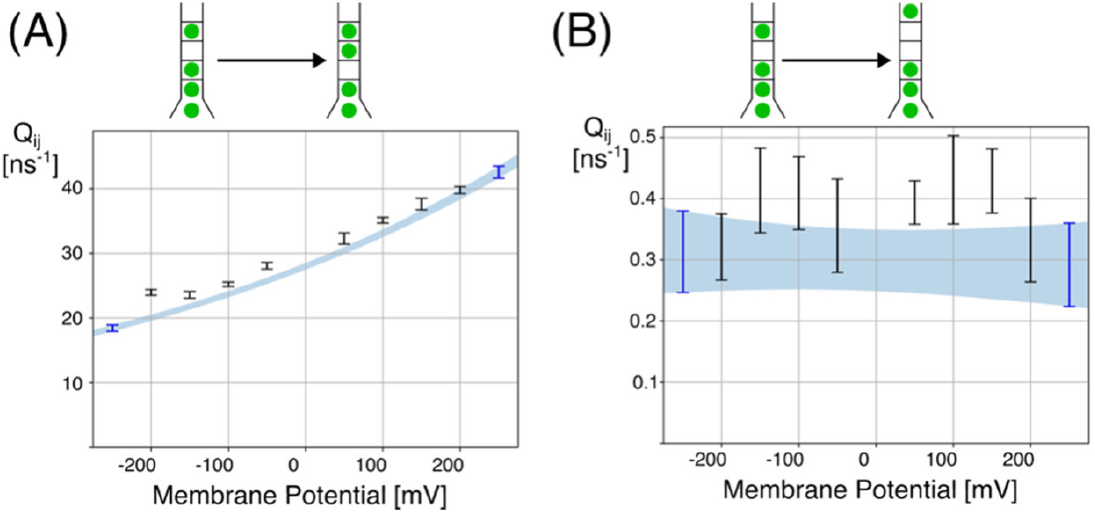
Voltage-dependence of rate constants in the MthK model. Two representative cases with respectively high and low voltage-dependence are shown. The entire rate matrix is reported in Figure S3. Blue-shaded area corresponds to the average estimated rate constants plus/minus one standard deviation, as computed by a bootstrap analysis. Error-bars are used to represents the values of rate constants computed directly from MD trajectories. Only the MD simulations are -250 mV and 250 mV (blue error-bars) were used for fitting the voltage-dependence of the rate constants. The schematic diagrams on the top represents the ion occupancy states in the terminal states. **(A)** Rate constant for the transition from a state with ions in C, S4, S3, S1 to a state with ions in C, S4, S2, S1. **(B)** Rate constant for the transitions from a state with ions in C, S4, S3, S1 to a state with ions in C, S4, S3, S0.

The channel current computed by equation (7) using the rate matrixes estimated by equation (9) is shown in Figure 3. The predicted current agrees with the values computed by counting the number of conduction events in MD trajectories. The estimated rate matrixes at different membrane voltages were also used to compute the steady-state probabilities of the 7 states included in the model. While the predictions do not always superimpose with the corresponding steady-state probabilities directly computed by MD, the qualitative trend is reproduced for all the 7 states and, in most cases, the deviations from MD data are minimal (Figure 4).

**Figure 3.**
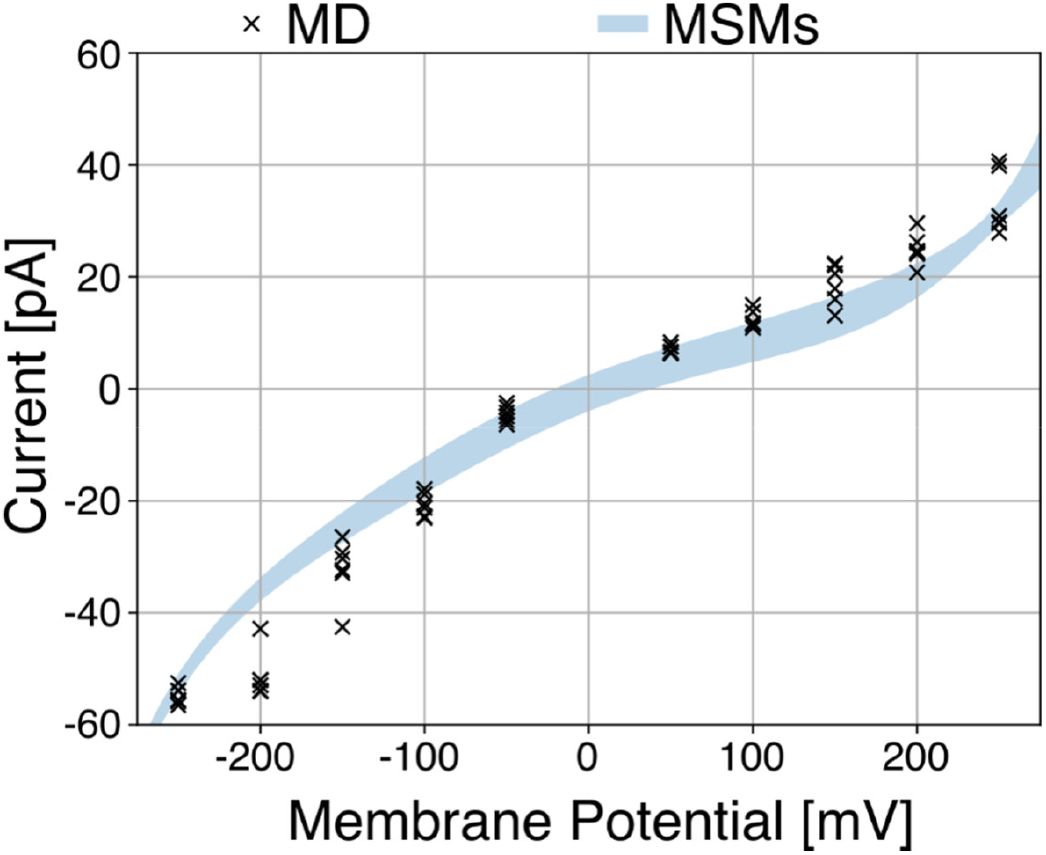
Current-voltage characteristic of the MthK model. The blue-shaded area corresponds to the average estimated current plus/minus one standard deviation, as computed by a bootstrap analysis. The currents computed by counting the number of conduction events in MD trajectories are shown as marks. Six independent trajectories are reported at each membrane potentials (Table S2).

**Figure 4.**
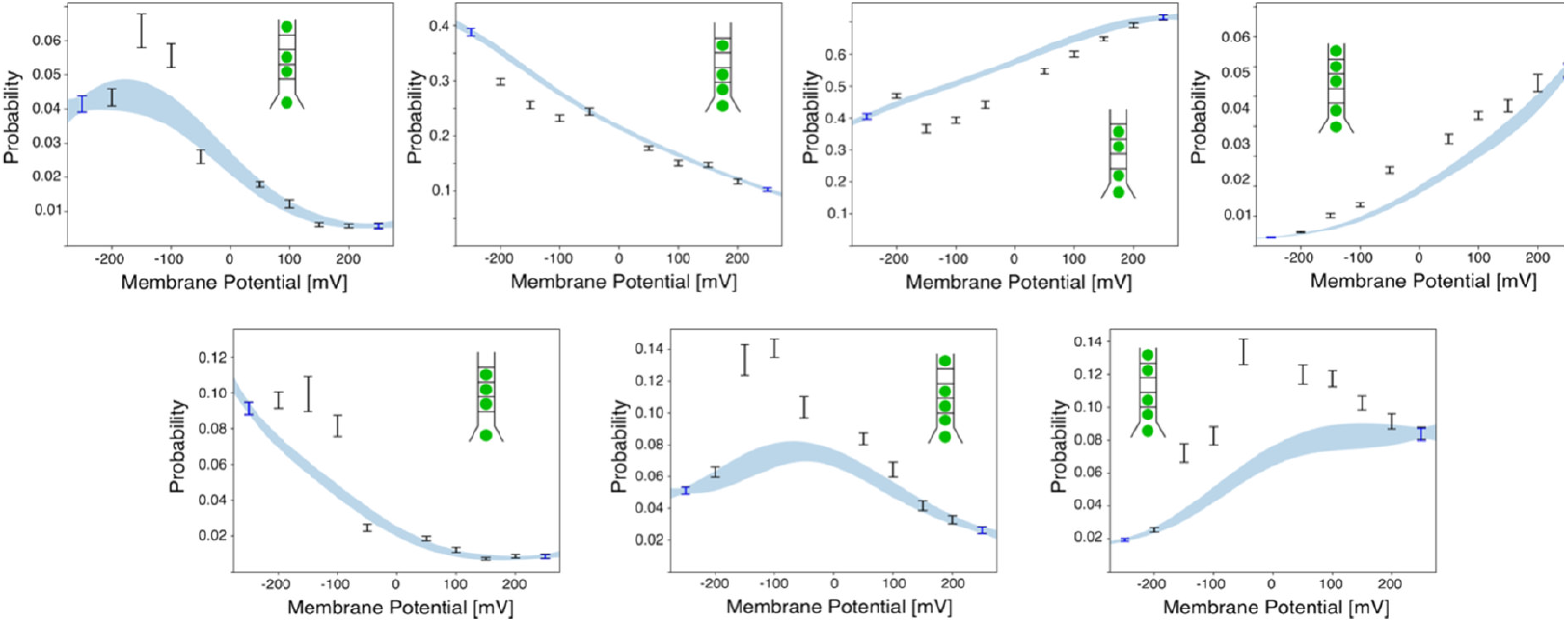
Steady-state probabilities in the MhtK model. The blue-shaded area corresponds to the average estimated steady-state probabilities plus/minus one standard deviation, as computed by a bootstrap analysis. Error-bars are used to represents the values computed from MSMs directly estimated from MD trajectories. Only the MD simulations at ±250 mV (blue error-bars) were used for fitting the voltage-dependence of the rate constants. The ion occupancy states are schematically depicted inside each plot.

### KcsA channel

The complete set of MD simulations for the KcsA model is summarized in Table S2. In MD trajectories obtained at negative membrane potentials, water molecules were observed to enter the central binding sites of the SF, S2 and S3 (Figure S4). These hydration events were effectively irreversible on the simulated timescale and became increasingly frequent as the membrane potential decreased. The long timescale associated with hydration of the SF preclude the reliable estimation of MSMs that include both de-hydrated and hydrated states. Specifically, water molecules entered the SF in 2 out of 4 trajectories at −100 mV, 4 out of 5 trajectories at −200 mV (with only one dehydration event observed), and in all 4 trajectories at −400 mV. At −400 mV, water molecules entered site S3 within less than 100 ns in three trajectories, resulting in only a few conduction events being observed before hydration occurred. In contrast, at −200 mV the dehydrated SF state persisted for several hundred nanoseconds, allowing a larger number of conduction events to be sampled. For these reasons, subsequent analyses were restricted to simulations performed at membrane potentials ≥ −200 mV and to trajectory segments in which sites S2 and S3 remained dehydrated, meaning that the model only describes conduction events with hard knock-on mechanism. The current–voltage characteristics of the KcsA model were therefore estimated using MD simulations at −200 mV and +400 mV.

The same procedure reported in previous section for the MthK model was applied to MD simulations of the KcsA model for selecting the sampling time (Figure S5), computing the rate matrixes (Figure S6), and computing the voltage dependence of the rate constants (Figure S7). There were 8 states with steady-state probability above 0.01 at membrane potential equal to - 200 mV or +400 mV, which cumulated respectively to 0.99 and 0.97 of the total probability (Table S4). The variability affecting the estimated rate constants is higher in the KcsA model compared to the MthK model, likely because of the lower number of conduction events sampled in KcsA, particularly in simulations with membrane potential equal to -200 mV. In accordance with observations in the MthK model, the rate constants associated with the movements of ions across the central binding sites of the SF exhibits a clear voltage-dependence (Figure 5A). Instead, the exit rate of an ion from S0 is practically unaffected by the membrane potential (Figure 5B).

**Figure 5.**
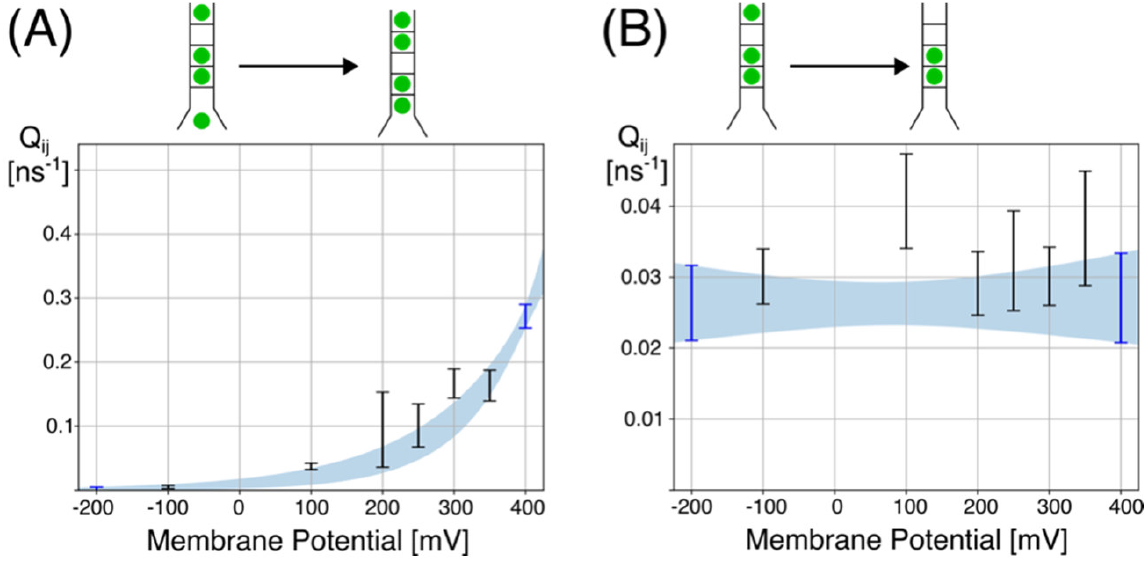
Voltage-dependence of rate constants in the KcsA model. Two representative cases with respectively high and low voltage-dependence are shown. The entire rate matrix is reported in Figure S7. The schematic diagrams on the top represents the ion occupancy in the terminal states. **(A)** Rate constant vs membrane potential for the transitions from a state with ions in C, S3, S2, S0 to a state with ions in S4, S3, S1, S0. **(B)** Rate constant vs membrane potential for the transitions from a state with ions in S3, S2, S0 to a state with ions in S3, S2.

The estimated current-voltage characteristic agrees with the currents computed by counting the number of conduction events in MD trajectories (Figure 6). In detail, the conductance at 100 mV is 4.3±2.2 pS when computed directly by MD, and 8.0±4.6 pS when estimated from the voltage-dependency of the rate matrix. As a comparison, the conductance estimated by MD (or MSMs) at +400 mV is 21.8±2.5 pS. The predictions of the model agrees with direct data from MD simulations also with respect to the steady-state probabilities (Figure 7). The model reproduces correctly also non-monotonic behaviours of the steady-state probabilities as a function of the membrane voltage. For instance, the probability of the state with ions in C, S3,S2 peaks at a membrane potential close to 100 mV, and the presence of this maximum is correctly identified by the model fitted using data at -200 mV and +400 mV.

**Figure 6.**
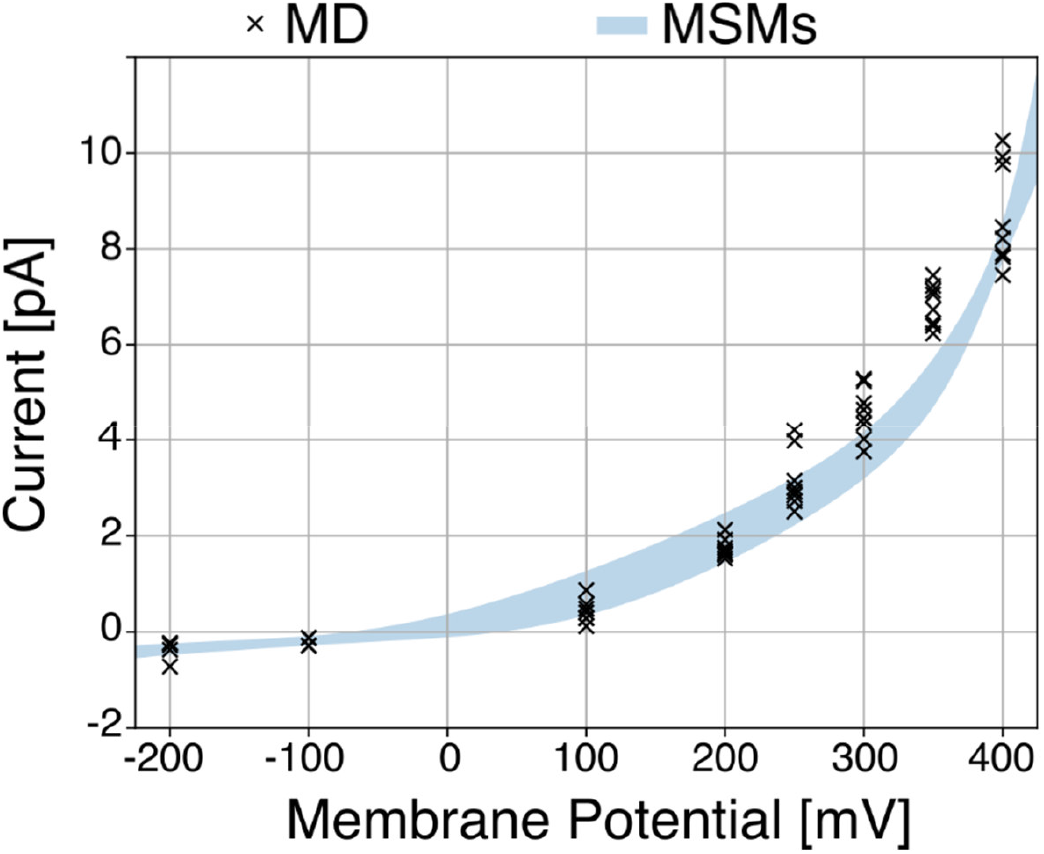
Current-voltage characteristic of the KcsA model. The blue-shaded area corresponds to the average estimated current plus/minus one standard deviation, as computed by a bootstrap analysis. The currents computed by counting the number of conduction events in MD trajectories are shown as marks (Table S2).

**Figure 7.**
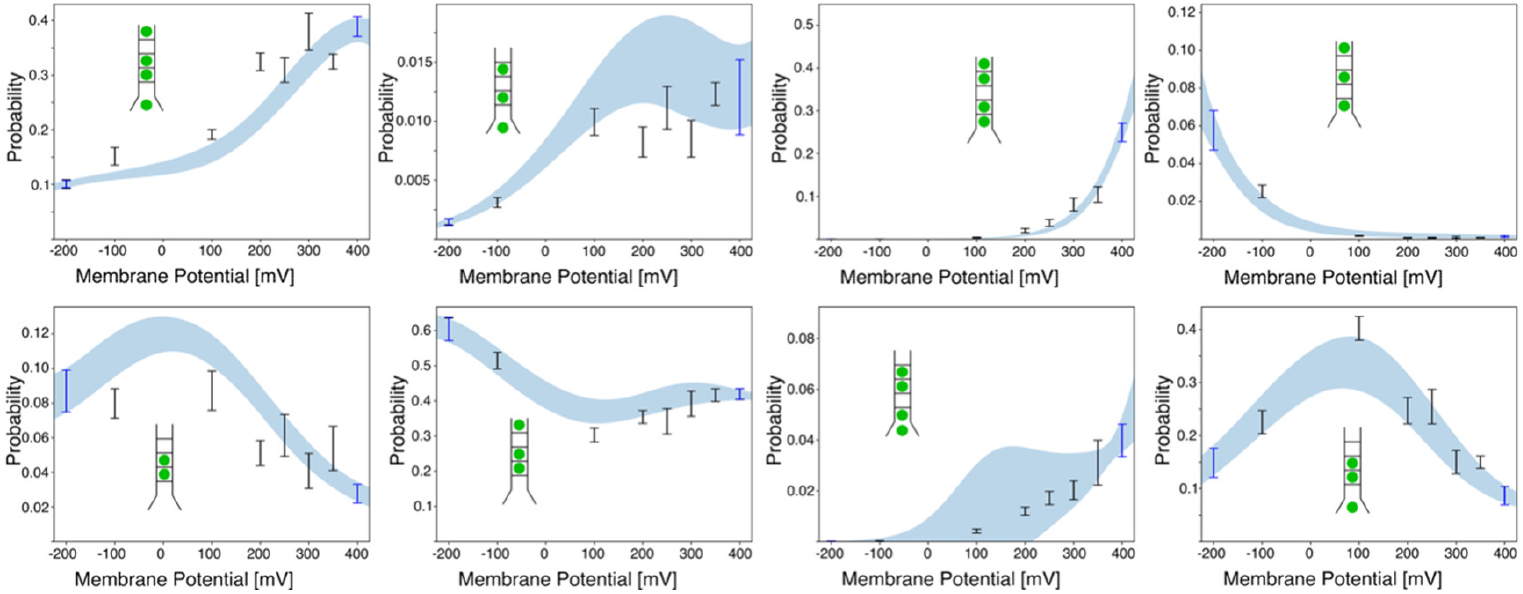
Steady-state probabilities in the KcsA model. The blue-shaded area corresponds to the average estimated steady-state probabilities plus/minus one standard deviation, as computed by a bootstrap analysis. Error-bars are used to represents the values computed from MSMs directly estimated from MD trajectories. Only the MD simulations at -200 mV and 400 mV (blue error-bars) were used for fitting the voltage-dependence of the rate constants. The ion occupancy states are schematically depicted inside each plot.

## Discussion

MSMs are emerging as a powerful strategy for analysing ion conduction in potassium channels starting from MD simulations (Harrigan *et al*., 2017)(Domene *et al*., 2021)(Kei & de Groot, 2023), or by combining atomic-level description with experimental data (Weckel-dahman *et al*., 2025). Recently, we proposed a method to compute currents across an ion channel directly from the transition or rate matrix of an MSM describing ion motion, in combination with a flux matrix that encodes the fractional charge transported through the channel for each pairwise transition (Catacuzzeno *et al*., 2024). Currents estimated using this approach show good agreement with direct estimates obtained from MD simulations. In the present study, we exploit this capability to compute channel currents directly from MSMs and develop a method to estimate the current-voltage characteristics of ion channels using MD simulations performed at only a limited number of membrane potentials. The method was validated by comparing its predictions with corresponding quantities computed from MD simulations at the same membrane potentials. Both the channel currents and the steady-state probabilities of the microscopic channel states predicted by our modelling approach were in satisfactory agreement with MD estimates.

A necessary condition to adopt the proposed method is the ability to estimate MSMs of ion conduction events from MD trajectories obtained at different membrane potentials. In addition, the same set of states, albeit with potentially very different probabilities, must be observed in simulations performed at all the membrane potentials used to fit the model. These requirements did not pose any difficulty when testing the proposed approach using the MthK model system. Owing to its high conductivity, MSMs of ion conduction could be readily constructed from MD simulations, and the same set of core states was observed at the two membrane potentials used for the fitting procedure. The situation was more complex for the KcsA model system. In this case, simulations performed at negative membrane potentials exhibited long-lasting transitions toward a hydrated state of the SF. The observation of these transitions is particularly relevant in the context of the ongoing discussion about soft and hard knock-on conduction mechanisms (Bosio *et al*., 2026), as it demonstrates that the presence of water molecules within the SF depends on the simulation conditions. However, addressing this mechanistic issue lies beyond the scope of the present study, which is focused on developing a method to estimate current-voltage characteristics from a limited number of MD simulations. Accordingly, our analysis was restricted to segments of the trajectories in which conduction occurs via the hard knock-on mechanism. Owing to the long timescales associated with hydration and dehydration of the SF in the KcsA model, it is not feasible to construct unified MSMs that simultaneously account for both hard and soft knock-on conduction. The resulting current-voltage characteristics should therefore be interpreted as representative only of channels conducting via the hard knock-on mechanism, which nevertheless constitutes the overwhelmingly dominant conduction mode in the KcsA model considered here.

The ability of our method to predict the ionic current at membrane potentials not directly tested in MD simulation relays on the assumption that the rate constants connecting the different states of the MSM are exponentially related to the applied membrane potential. As first shown by Woodhull, Hille and Schwartz in the case of ion channel permeation (Woodhull, 1973)(Hille & Schwarz, 1978), an exponential relationship between the kinetic rate constants and the applied membrane potential is expected when ions move across substantially high energy barriers present along the permeation pathways and separating stable ion binding sites. Under these conditions the ion diffusion process can be approximated with very fast transitions between adjacent binding sites, with kinetic rate constants exponentially related to the barrier energy height (Kramers, 1940)(Cooper *et al*., 1988). If an electric potential drop exists between the two binding sites, then the peak energy will have an electrostatic component directly proportional of the percentage of voltage applied to the membrane dropping between the energy well and the energy peak. Notably, numerous experimental studies on potassium channel permeation confirm the validity of this assumption along the permeation pathway, by showing that the movement of ions inside the channel pores behaves according to rate constants with an exponential voltage relationship, see for instance (Miller *et al*., 1987)(Nimigean & Miller, 2002)(Neyton & Miller, 1988). In addition, our own MD simulations and MSM approach shows that the changes of the kinetic rate constants with the applied potential are well described by exponential relationships, as theoretically expected.

Besides predicting the current at applied potentials not directly tested with MD simulations, the method presented here is also able to define in detail the voltage-dependence of the permeation mechanism. For example, in MthK channels the steady-state occupancy of the relevant configuration shows that states with three binding sites are favoured by membrane hyperpolarization, while states with 4 binding sites occupied by potassium ions are favoured by depolarized potentials. A similar trend of increasing number of occupied sites with depolarization is present also in KcsA channels, although in this case a significant lower number is evident as compared to MthK. Interestingly, the method proposed appears able to predict the voltage dependent changes in steady state occupancy, with maxima or wells, present at intermediate voltages not used as input, quite well reproduced.

Provided that the conditions outlined in the previous paragraphs are satisfied, the method proposed here offers a valuable framework for bridging MD simulations and experimental measurements. Due to their high computational cost, MD simulations of ion conduction are typically performed at membrane potentials substantially higher than physiological values, in order to enhance sampling of conduction events. The approach introduced here enables the use of MD simulations carried out at elevated membrane potentials while still allowing the estimation of channel conductance under physiological conditions. As accessible MD simulation timescales have increased, quantitative discrepancies between MD-based estimates of ion channel conductance and experimental observations have become more apparent. Considerable effort is currently devoted to refining force fields to improve agreement with experimental data. By enabling more direct and systematic comparisons between simulation-based predictions and experimental measurements, our method provides a useful tool to support and guide these ongoing modelling efforts.

## Author Contributions

The manuscript was written through contributions of all authors. All authors have given approval to the final version of the manuscript.

## Acknowledgments

We acknowledge CINECA for awarding access to computational resources through the ISCRA Initiative (grant numbers HP10B597KB and HP10B5IPGG).

## Notes

Discretized MD trajectories, and the python code used for the analyses of the MSM are available at the github repository: https://github.com/sfurini/current_voltage_md_msm

## Abbreviations

MD: Molecular Dynamics
SF: Selectivity Filter
MSM: Markov State Model.

