## Supplementary Information for "Current-voltage characteristics of K+ channels estimated by MD simulations and Markov State Models"

| **Membrane Potential [mV]** | **Simulation Time [µs]** | **Number of conduction events** |
|---|---|---|
| -250 | 6 x 0.5 | -164, -173, -168, -176, -174, -168 |
| -200 | 6 x 0.5 | -162, -165, -168, -162, -169, -134 |
| -150 | 6 x 0.5 | -102, -91, -101, -133, -83, -95 |
| -100 | 6 x 0.5 | -59, -64, 56, 66, -71, -72 |
| -50 | 6 x 0.5 | -13, -18, -15, -10, 20, -8 |
| 50 | 6 x 0.5 | 20, 26, 21, 20, 24, 23 |
| 100 | 6 x 0.5 | 36, 34, 43, 47, 37, 36 |
| 150 | 6 x 0.5 | 50, 41, 70, 64, 56, 69 |
| 200 | 6 x 0.5 | 92, 75, 77, 65, 82, 76 |
| 250 | 6 x 0.5 | 92, 127, 96, 87, 93, 124 |

**Table S1. Molecular Dynamics simulations of the MthK model.** Number of conduction events are reported separately for each independent replica at each membrane potential (minus sign indicates inward conduction events).

| Membrane Potential [mV] | Simulation Time [µs] | Number of conduction events |
|---|---|---|
| -400 | 1.0 (<0.1), 1.0 (<0.1), 1.0 (<0.1), 1.0 (0.5) | -2, -1, -1, -3 |
| -200 | 5.0 (3.2), 5.0 (2.0), 5.0 (0.5), 5.0, 5.0 (1.75) | -5, -9, -1, -11, -2 |
| -100 | 5.0, 5.0, 5.0 (2.0), 5.0 (3.2) | -4, -9, -1, -5 |
| 100 | 7 x 4.0 | 9, 19, 1, 12, 5, 7, 7 |
| 200 | 8 x 4.0 | 48, 40, 41, 40, 53, 44, 42, 38 |
| 250 | 8 x 2.0 | 34, 39, 37, 31, 35, 53, 36, 50 |
| 300 | 8 x 1.0 | 33, 28, 33, 30, 29, 23, 25, 27 |
| 350 | 8 x 1.0 | 46, 40, 42, 39, 45, 44, 40, 44 |
| 400 | 8 x 1.0 | 47, 62, 51, 49, 61, 64, 53, 49 |

**Table S2. Molecular Dynamics simulations of the KcsA model.** Number of conduction events are reported separately for each independent replica at each membrane potential (minus sign indicates inward conduction events). In trajectories where water molecules were observed to enter into binding sites S2 or S3, the simulation time in the absence of water molecules is reported in brackets.

| State | Steady-state probability | |
|---|---|---|
| | **-250 mV** | **250 mV** |
| C, S3, S2, S0 | 0.04 | 0.006 |
| C, S4, S3, S1 | 0.38 | 0.10 |
| C, S4, S2, S1 | 0.40 | 0.71 |
| C, S4, S2, S1, S0 | 0.003 | 0.06 |
| C, S3, S2, S1 | 0.09 | 0.008 |
| C, S4, S3, S2, S0 | 0.05 | 0.03 |
| C, S4, S3, S1, S0 | 0.02 | 0.08 |

**Table S3. Steady-state probabilities in the MthK model.** Only states with probability above 1% in simulations at -250 mV or +250 mV are reported.

| State | Steady-state probability | |
|---|---|---|
| | -200 mV | 400 mV |
| C, S3, S2, S0 | 0.10 | 0.36 |
| C, S3, S1 | 0.001 | 0.01 |
| S4, S3, S1, S0 | $10^{-6}$ | 0.02 |
| S4, S2, S0 | 0.05 | 0.001 |
| S3, S2 | 0.09 | 0.03 |
| S3, S2, S0 | 0.60 | 0.41 |
| C, S4, S2, S1 | $10^{-4}$ | 0.04 |
| C, S3, S2 | 0.15 | 0.09 |

**Table S4. Steady-state probabilities in the KcsA model.** Only states with probability above 1% in simulations at -200 mV or +400 mV are reported.

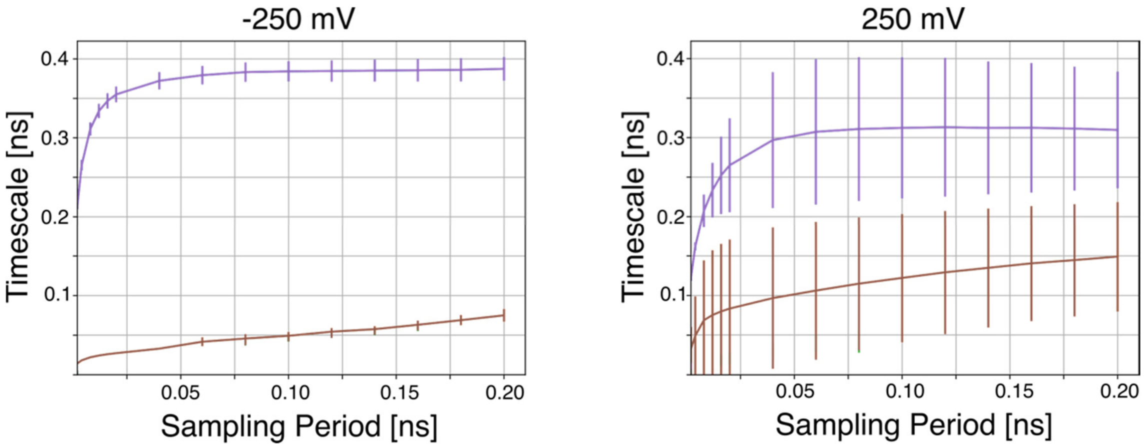


**Figure S1. Relaxation times of the MSMs for the MthK model.** Data are shown for simulations at -250 mV and +250 mV.

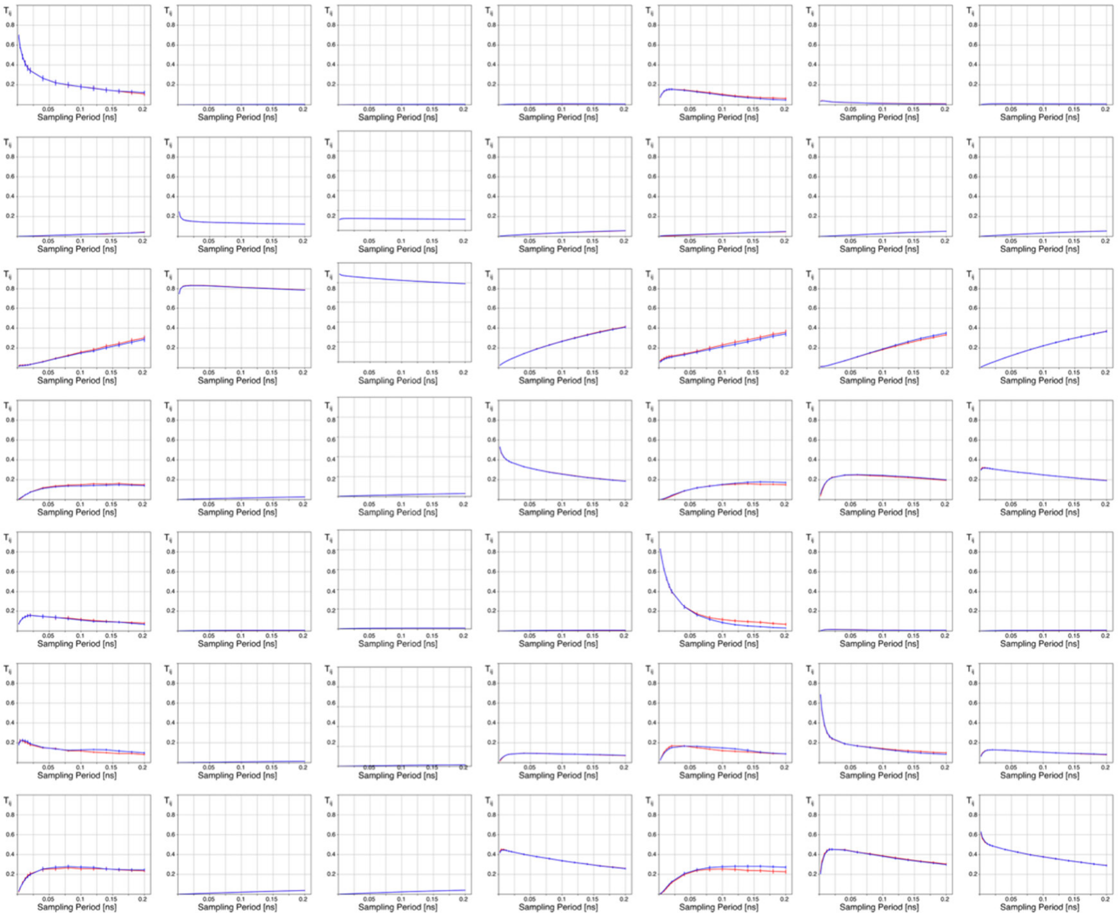


**Figure S2. Transition matrixes estimated by MD and computed from rate matrixes.** Plots of the transition probabilities vs the lagtime for all elements of the transition matrix. Red lines are used for the elements of the transition matrix as estimated directly by MD simulations. Blue lines are lines are used for the elements of the transition matrix estimated from the rate matrix by equation (6). Data refers to MD simulations of the MthK model at +250 mV. States are ordered as in Table S3.

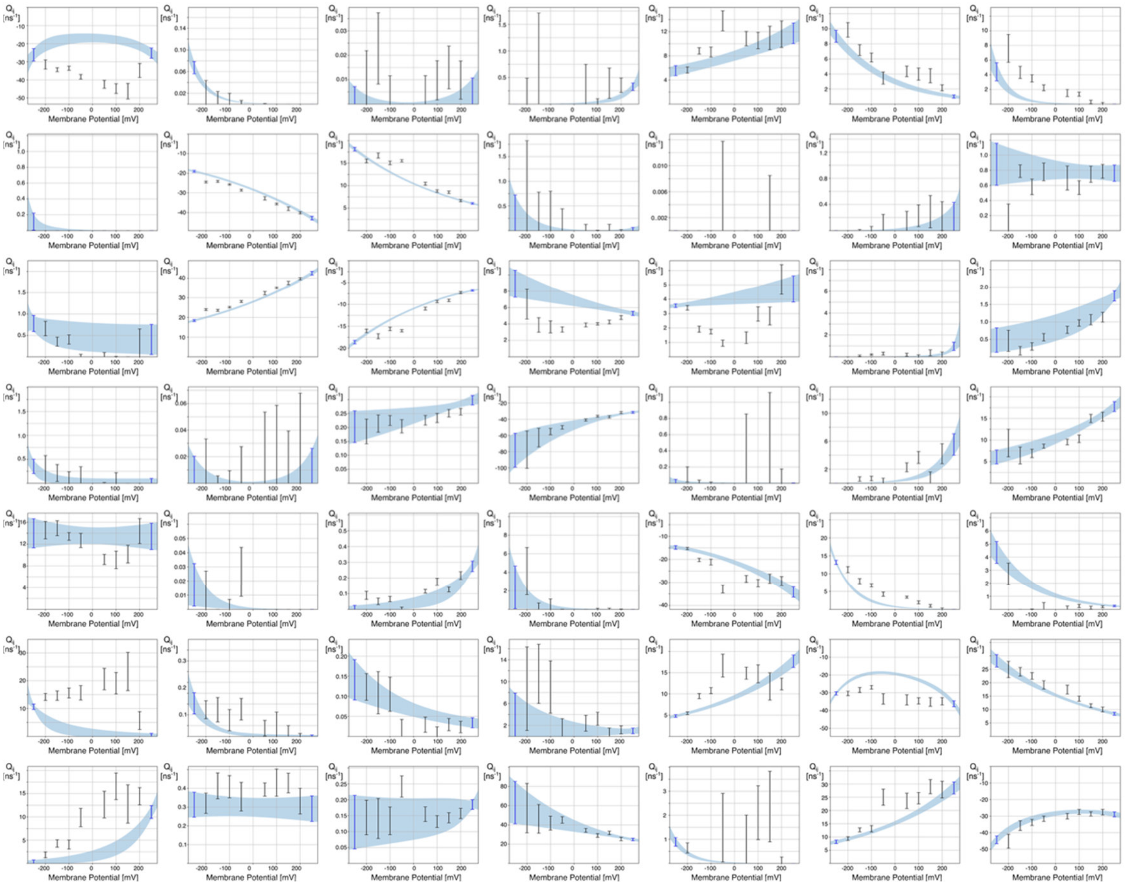


**Figure S3. Voltage dependence of the rate constants in the MthK model.** Blue-shaded area corresponds to the average estimated rate constants plus/minus one standard deviation, as computed by a bootstrap analysis. Error-bars are used to represents the values of rate constants directly estimated from MD trajectories. Only the MD simulations at -250 mV and 250 mV (blue error-bars) were used for fitting the voltage-dependence of the rate constants. States are ordered as in Table S3.

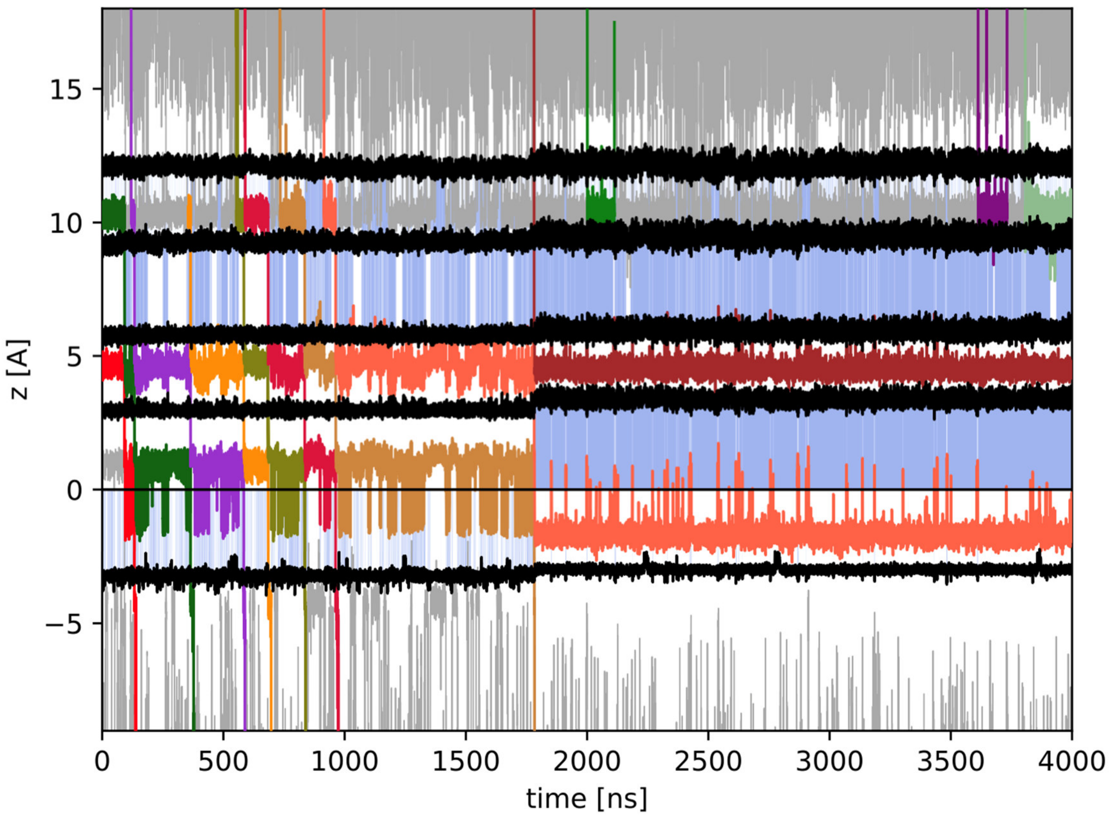


**Figure S4. MD trajectory of the KcsA model at -200 mV.** Black lines corresponds to the boundaries of binding sites S4-S0, from the bottom. Coloured lines are used for ions inside the SF, grey lines for ions outside the SF. Blue-shading is used for binding sites occupied by water molecules.

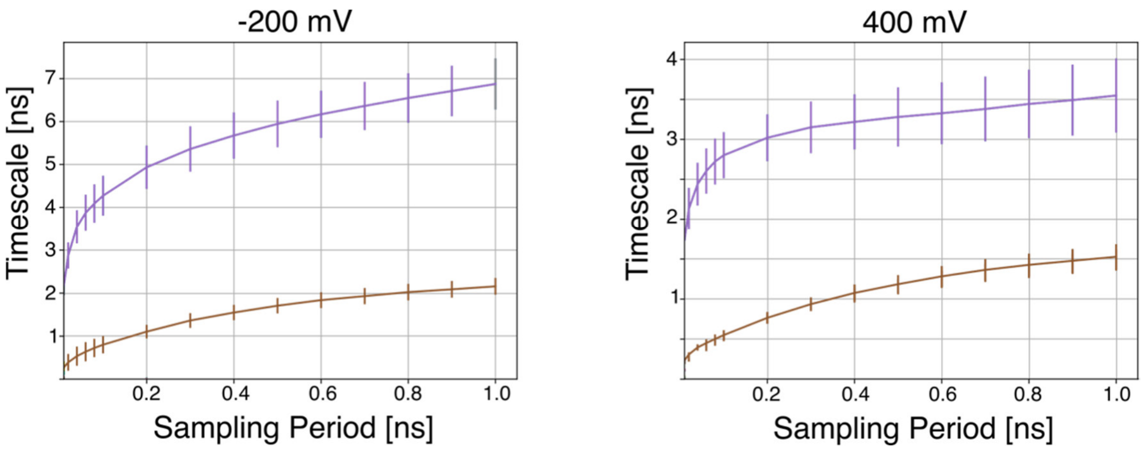


**Figure S5. Relaxation times of the MSMs for the KcsA model.** Data are shown for simulations at -200 mV and +400 mV.

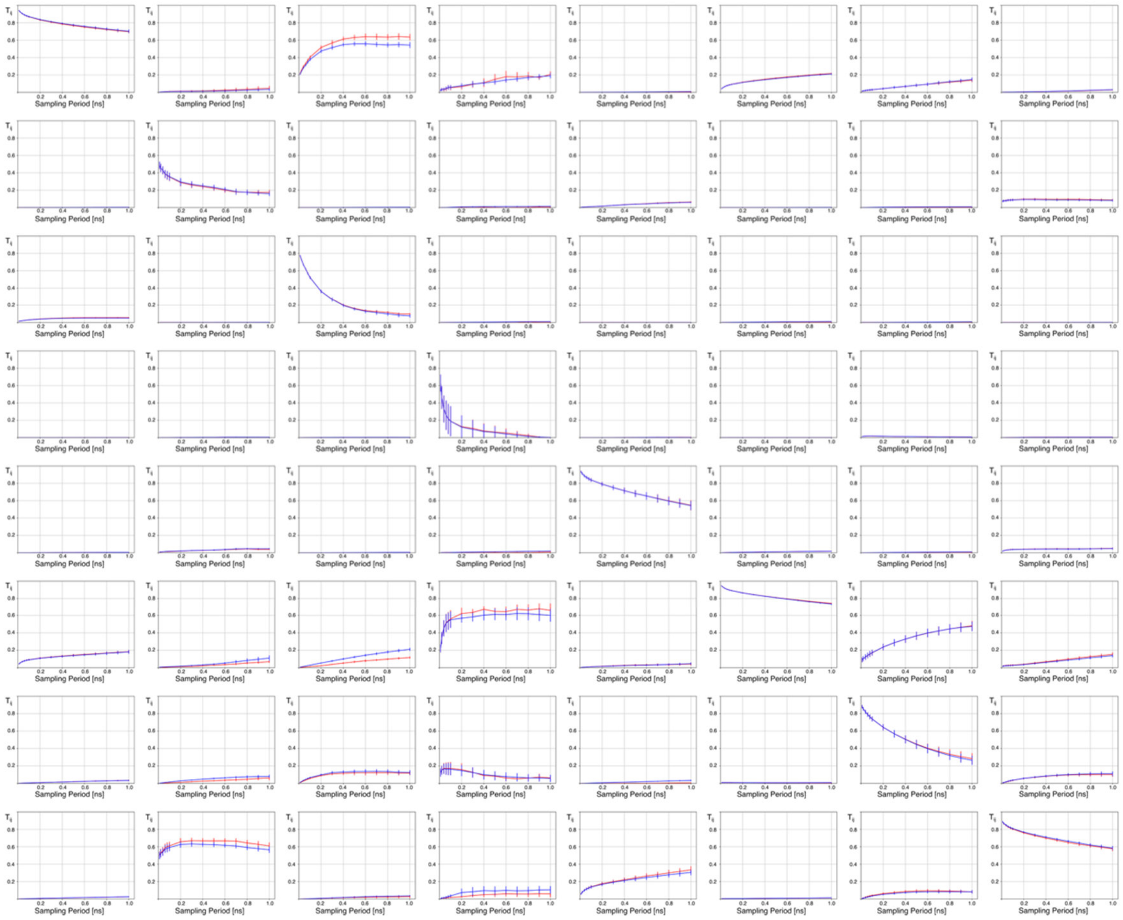

**Figure S6. Transition matrixes estimated by MD and computed from rate matrixes.** Red lines are used for the elements of the transition matrix as estimated directly by MD simulations. Blue lines are used for the elements of the transition matrix estimated from the rate matrix by equation (6). Data refers to MD simulations of the KcsA model at +400 mV. States are ordered as in Table S4.

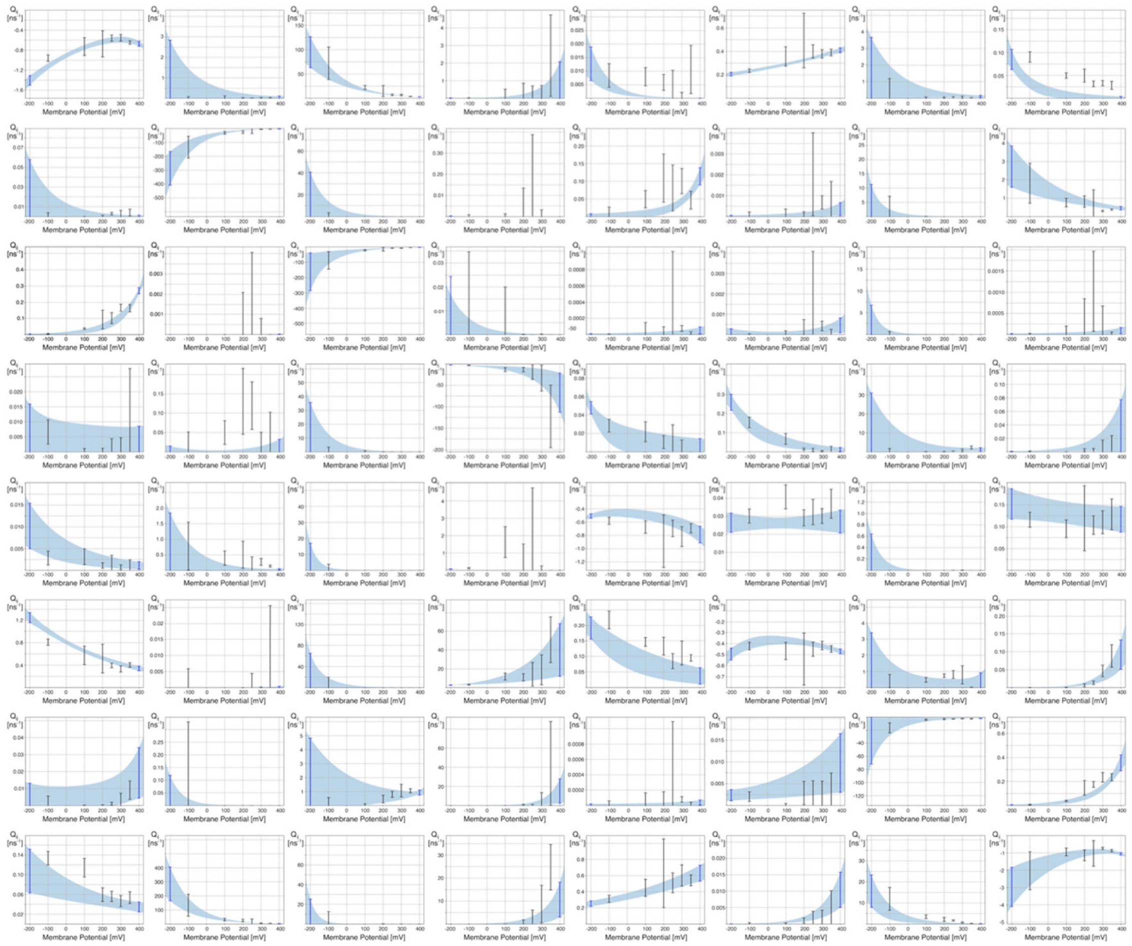

**Figure S7. Voltage dependence of the rate constants in the KcsA model.** Blue-shading corresponds to the average estimated rate constants plus/minus one standard deviation, as computed by a bootstrap analysis. Error-bars are used to represents the values of rate constants directly estimated from MD trajectories. Only the MD simulations at -200 mV and 400 mV (blue error-bars) were used for fitting the voltage-dependence of the rate constants. States are ordered as in Table S3.
